# Fractal Measures as Predictors of Histopathological Complexity in Breast Carcinoma Mammograms

**DOI:** 10.1101/2025.04.10.648305

**Authors:** Abhijeet Das, Ramray Bhat, Mohit Kumar Jolly

## Abstract

Breast carcinoma remains the most commonly diagnosed malignancy and a leading cause of cancer-related mortality among women worldwide. While mammography is the gold standard for early detection, challenges such as high breast density often obscure malignancies, reducing diagnostic sensitivity. Conventional parenchymal texture analysis methods have limitations due to struggles with spatial interpretation and noise sensitivity. This study investigates the efficacy of fractal-based global texture features for distinguishing between malignant and normal mammograms and assessing their potential for molecular subtype differentiation. Digital mammograms were analyzed using standardized preprocessing techniques, and fractal measures were computed to capture complexity and connectivity properties within breast tissue structures. Fractal dimension, multifractality strength, and succolarity reservoir were found to effectively characterize specific features of mammographic texture; however, their incorporation into machine learning models yielded moderate discriminatory performance between categories. We introduced succolarity reservoir as a novel parameter accounting for tissues’ latent connectivity. In addition, while succolarity reservoir exhibits potential for differentiating Luminal B from other molecular subtypes, its overall discriminative power remains limited. This proof-of-concept study underscores the potential of fractal-based texture analysis as a non-invasive biomarker in breast carcinoma diagnosis.

## 1. Introduction

Breast carcinoma (BC) is the most frequently reported form of cancer and the second leading cause of cancer-related mortality in women (1). Early diagnosis is crucial for guiding treatment strategies, improving prognosis, and reducing mortality rates. Mammography remains the gold standard for screening and early detection/diagnosis of BC, offering a non-invasive approach to detecting abnormalities (2).

The mammographic density of tissues contributing to the overall density of breasts is a critical biomarker and independent predictor that indicates the possibility of developing BC. Women with high breast density can have a 4 to 6-fold higher chance of developing BC than women with low breast density (3). In addition, high breast density, caused by a higher proportion of fibro-glandular tissues (consisting of epithelial and stromal cells), obscures malignancies by causing a masking effect on lesions with masses or architectural distortions, subsequently reducing sensitivity and increasing the likelihood of false negatives in screening mammography (4), particularly in women less than 50 years of age (5). The issue is also relevant in terms of demography, especially concerning women of Asian ethnicity, who tend to have a more prevalent average breast density than their Western counterparts (6), further complicating the distinction between normal and malignant tissues. Notably, the mammographic percent density of Chinese women was significantly higher than that of women from Malaysia and India, even after adjusting for the body mass index, age, menopausal status, etc. (7).

Texture provides us with a visual cue to the repetition of patterns in an image and can be informally defined as a structure composed of a large number of more or less ordered similar patterns. In image analysis, a texture pattern refers to the spatial arrangement of pixels that is insufficiently defined by regional intensity or color alone (8). In the context of mammogram images, texture is defined as the spatial relationship and variations in grey-scale values among pixels within a specified region of interest (ROI) of the breast. In a biophysical sense, it characterizes the spatial distribution and heterogeneity of parenchymal tissue patterns, reflecting the proportion of glandular tissue to fatty tissue. This biophysical characterization reflects a correlation between breast density and parenchymal texture, as it highlights the distribution of high (low) density and textured glandular (fatty) breast tissues across the mammographic image.

Mammographic texture is mainly studied using three approaches, viz., statistics-based, model-based, and transform-based (9). Among them, statistics-based methods are the most commonly utilized, including analysis of texture using first-order statistics (histogram), second-order statistics (grey-level co-occurrence matrix), and higher-order statistics (run length matrix, size zone matrix). On the other hand, model-based approaches like fractal geometry and transform-based approaches like the Fourier transform, wavelet and Gabor filters, and Laplacian transform of Gaussian filter, respectively, are scarcely utilized (9). Nevertheless, the statistics-based methods have certain limitations, such as ignoring the spatial arrangement of pixels, dependency on the choice of displacement vector and angle, and consideration of fixed distance between pixels (9,10). Additionally, the transform-based methods like (i) Fourier transform-based approaches represent textures in the frequency domain and capture periodic texture patterns but do not preserve the locally irregular patterns and direct interpretation of spatial localization of textures, respectively, (ii) Wavelet transform-based approaches require choice of the mother wavelets (e.g., Haar, Daubechies) and decomposition level, which subsequently require careful optimization and are not rotationally or translationally invariant thus affecting the outcomes (11), and (iii) Laplace transform-based methods can amplify noise owing to emphasis on high-frequency components, especially in low-contrast mammograms (9). A common limitation of most studies using the texture analysis approaches is their limited interpretation from a biological and/or biophysical viewpoint (9). However, model-based approaches like mono- and multi-fractal analysis of mammograms can be useful because they could capture statistically self-similar and heterogeneous patterns in breast tissue, remain invariant across different resolutions or scales, and are less sensitive to noise (12).

Texture analysis in mammogram images has been widely explored as a potential computer-aided diagnostic method to enhance BC diagnosis, particularly for differentiating tissue composition or detecting malignancies. However, previous studies have primarily relied on statistics-, transform-, graphical-, and lattice-based approaches (13–23). These methods have demonstrated considerable success, achieving high classification accuracies, particularly when coupled with machine learning algorithms (14,15,19). Nonetheless, their ability to provide a biophysically meaningful interpretation of tissue heterogeneity and malignancy-related structural complexity and connectivity remains limited. Although deep-learning approaches to extract high-dimensional feature representations from mammograms, such as convolutional neural networks (CNNs) trained on pixel-level data (24–26), have shown promise, they typically require large annotated datasets for robust training, posing a challenge owing to the limited availability of required datasets. Furthermore, deep-learning methods often lack interpretability, making it difficult to link extracted features to meaningful biophysical and histological properties of breast tissues.

To address these challenges, recently, mono- and multi-fractal analysis has been proposed as an alternative approach to capture self-similarity, heterogeneity, and complexity in mammographic textures (19,27,28). Given their ability to quantify structural or architectural organization in tissue patterns at multiple spatial scales, fractal-based methods can provide insights into the complexity of breast tissue that conventional statistical measures may not fully capture, and can allow differential detection or diagnosis of BC along with classification frameworks (5, 27, 29, 30-34). This approach is based on the premise that tumor boundaries exhibit greater irregularity than normal tissues. However, contour-based analysis possesses limitations, as tumor margins in mammograms are often poorly defined owing to low contrast, noise, or overlapping structures. Also, the datasets utilized in these studies have been from the established databases from the western parts and are often limited in size.

BC is also classified into 4 subtypes according to gene expression profiles, where each subtype corresponds to varied prognosis, risk of progression, response to treatment, and survival outcomes (35). These 4 surrogate subtypes, i.e., luminal A, luminal B, human epidermal growth factor receptor 2 (HER2)-enriched, and triple-negative, are based on semiquantitative immunohistochemistry (IHC) scoring of estrogen receptor (ER), progesterone receptor (PR), and *in-situ* hybridization test for HER2 overexpression (36). While IHC and genomic profiling remain the standard protocol for subtype classification, these methods are invasive, costly, and prone to sampling bias due to intratumoral heterogeneity. Consequently, non-invasive imaging biomarkers that reflect molecular diversity have emerged as a critical frontier in precision oncology.

This study evaluates the effectiveness of mono- and multi-fractal measures, including fractal dimension, lacunarity, succolarity reservoir, Renyi dimensions, and multifractality strength in distinguishing normal from malignant mammograms and among molecular subtypes, based on global parenchymal texture analysis and their biophysical interpretation. Sections 2–5 cover the dataset, preprocessing, fractal feature computation, statistical and machine learning methods, results, biophysical interpretation, methodological insights, and conclusions with directions for future research.

## 2. Methodology

### 2.1 Dataset and Preprocessing

This study primarily focuses on digital mammograms with a Mediolateral Oblique (MLO) view and standard dimensions of 1914 × 2294 pixels from the Chinese Mammography Database (CMMD), specifically CMMD2 (37), containing 1498 mammograms from 749 cases of women with a mean age of 49.82 years and a median age of 49 years. It also contains information regarding the molecular subtypes evaluated using the Core Needle Biopsy technique. However, since no information was provided regarding the ROIs in the original dataset [https://www.cancerimagingarchive.net/collection/cmmd/], we utilized the TOMPEI-CMMD dataset, containing information about the ROIs in JSON format for the malignant cases, assessed by a practicing radiologist with 20+ years of experience. Following this, mammograms with undetectable lesions and insufficient image quality were excluded from the dataset. The dataset can be accessed from the given link [https://www.cancerimagingarchive.net/analysis-result/tompei-cmmd/]. Subsequently, we have analyzed 1778 ROIs belonging to the normal and malignant categories from 671 cases.

Mammography images are known to contain contrast and background noise in varying degrees and, consequently, necessitate preprocessing steps for even distribution of contrast and suppressed noise in the image (38). In this context, we employed the bottom and top hat operations for accentuating both bright and dark regions in the mammogram and improving the visibility of fine structures. The top (bottom) hat transformation enhances the high- (low-) intensity regions in an image by subtracting the result of a morphological opening (closing) from the original image. Opening is a fundamental operation in mathematical morphology that removes small bright structures or noise while preserving the overall shape of the original object using two sequential operations, namely, erosion and dilation. Here, erosion shrinks the bright regions by removing pixels not fitting within the structuring element, and deletion expands the remaining bright regions back to their original size. Similarly, closing removes small dark structures and smooths objects’ boundaries. Here, erosion expands the bright regions by filling gaps within the structuring element, and deletion shrinks the expanded regions back to their original shape while keeping the filled structures. Mathematically, the utilized top and bottom hat operations are given by Eq. (1) and Eq. (2), respectively.

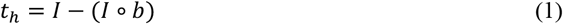

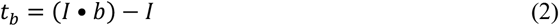

Here, *t*_*h*_ and *t*_*b*_ represents the Top and Bottom hat operation, and ∘ and • represents the morphological opening and closing operators, respectively. The original image is represented by *I* and *b* represents the structuring element (rectangular kernel) of size 15 × 15. The size was implemented to enhance both the small- and large-scale features, like microcalcification and mass in the mammograms.

The ROIs were extracted using the Fiji/ImageJ2 software and saved in TIF format. Since the origin of the coordinate system was not provided in the original work, we assumed it to be according to the standard norm, i.e., the top-left corner. Also, for the normal category and in consideration of the breast laterality (left or right), ROIs were determined and extracted using the information for the malignant category, with a transformation in the *x*-coordinate using Eq. (3)

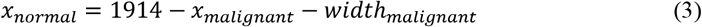

Many of the image-processing techniques, like fractal feature extraction, classification, and machine and deep learning models, require a fixed input size. In the analyzed dataset, ROIs had varying dimensions, and direct resizing of an image can alter the original aspect ratio, leading to distortion of structural features, introducing edge artifacts or artificial contrast shifts, and introducing bias in machine learning models. In this regard, we standardized the dimensions of all the ROI images to 256 × 256 pixels while maintaining a constant aspect ratio by resizing the original image using the Lánczos interpolation for rescaling, centering the image, and then utilizing neutral grey (intensity = 127) padding equally on all sides. The grey padding was used to minimize the contrast difference with the ROI images and avoid the introduction of high-contrast edges, unlike those created by black (0) and white (255) padding.

### 2.2 Computation of Mono- and Multi-fractal Measures

The mono- and multi-fractal parameters were computed using the ComsystanJ plugin in Fiji (39). The multifractality strength was computed from the multifractal spectrum. The methods involved in determining specific parameters are discussed below.

#### 2.2.1 Fractal Dimension

Sarkar and Chaudhuri proposed the Differential Box-Counting (DBC) algorithm (40) for computing the fractal dimension of textured images, which is based on the three-dimensional representation of a two-dimensional grey-scale image, where the *x* and *y* coordinates correspond to the spatial coordinates (image pixels) and the *z* coordinate represents the grey-level intensity of each pixel. However, the approach assumes equal step sizes in the grey-level division and does not consider relative changes in the grey level. The mentioned may introduce errors in the estimation of textural complexity, especially in highly textured images like mammograms. Subsequently, Jin *et al*. (41) improved upon the DBC method and proposed the Relative Differential Box-Counting (RDBC) method for the computation of fractal dimension. Similar to the DBC method, the image is treated as a three-dimensional surface. Here, an image of size *M* × *M* is divided into grids of size ϵ × ϵ, where ϵ represents the scale factor. In this work, we varied ϵ from 2^1^ to 2^9^. However, instead of using the absolute box height difference, the method computes the relative difference for each grid using Eq. (4) (41).

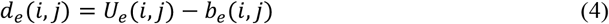

In Eq. (4), *U*_*e*_(*i, j*) and *b*_*e*_(*i, j*) represents the maximum and minimum grey levels in the grid, respectively. Hereafter, the number of occupied boxes in each grid is computed using Eq. (5).

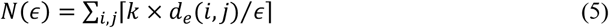

Here, *k* = *M*/*G* is the scaling factor in the *z*-axis and ⌈ ⌉ is the ceiling function. Finally, the fractal dimension of the textured image is computed from the best line fit of *N*(ϵ) and ϵ plot on a log-log scale.

#### 2.2.2 Lacunarity

Lacunarity is a scale-dependent measure of texture heterogeneity and clustering. While the fractal dimension provides information regarding self-similarity and complexity, lacunarity is used to acquire additional information regarding voids and structural variations in the texture. Also, since the raster-box method was used for computing the fractal dimension measure, we have adopted the same method for avoiding bias in feature comparison across categories and computed the lacunarity measure utilizing the methodology reported by Roy and Perfect (42). Non-overlapping boxes of size (ϵ × ϵ) was placed over an image with ϵ varied from 1 to 9 in powers of 2. Subsequently, lacunarity for each box size was computed using Eq. (6).

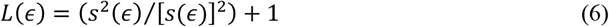

Here, *s*(ϵ) and *s*^2^(ϵ) represents the total mass (or pixel intensity) and squared mass of pixels within each box of size ϵ.

#### 2.2.3 Succolarity Reservoir

The conventional succolarity parameter measures the degree of percolation in specific directions; however, the succolarity reservoir provides a global overview of percolation capacity or connectivity behavior and is defined from Eq. (7) (43)

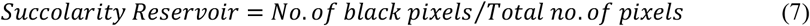

#### 2.2.4 Renyi Dimensions

The Renyi dimensions values were extracted from the computation of the Generalized dimensions (*D*_*q*_) which provides a spectrum of fractal measures by quantifying the mass distribution across scales, for describing the multiscale textural heterogeneity of an image. We adopted the methodology described by Ahammer *et al*. (44). Here, a mammogram image is divided into non-overlapping boxes of size (ϵ × ϵ), where ϵ varies from 1 to 9, and the number of pixels in each box is computed. The probability of mass *p*_*i*_(ϵ) inside each box *i* is computed using Eq. (8).

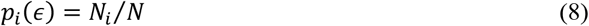

Here, *N*_*i*_ is the number of pixels in the *i*th box and *N* is the total number of pixels in the image. The generalized correlation integral is given by Eq. (9):

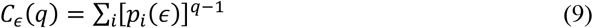

Where, *q* (mass exponent) controls the emphasis on different intensity regions in an image. The generalized dimensions are obtained from the scaling behavior of Eq. (9) by fitting a linear regression to the log-log plot of *C*_ϵ_(*q*) vs. ϵ.

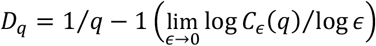

Subsequently, the Renyi dimensions, i.e., capacity, information, and correlation dimensions, were computed using 10(a), 10(b), and 10(c), respectively.

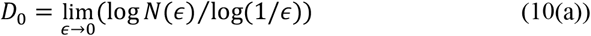

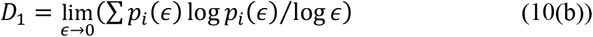

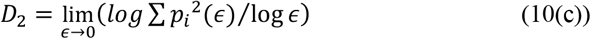

#### 2.2.5 Multifractality Strength

The multifractality strength is computed from the multifractal spectrum. The multifractal spectrum is plotted between two parameters, namely, the singularity strength (*α*(*q*)) and the fractal dimension of the set (*f*(*α*)), computed from Eq. (11) using the finite difference method and Eq. (12), respectively (45,46).

### 2.3 Statistical Analysis

A comprehensive statistical analysis was performed to assess the ability of mono- and multi-fractal texture features to differentiate between normal and malignant categories and across different breast cancer subtypes. A *p*-value < 0.05 was considered statistically significant.

#### 2.3.1 Descriptive Statistics and Normality Assessment

For each computed fractal feature, the descriptive statistical measure (mean, standard deviation, median, histograms, box plots, and violin plots) was computed separately for Normal vs. Malignant categories and across subtypes. Violin plots were used for subtypes since they consist of multiple groups, thus allowing better visualization of distribution shape and density, respectively. The Shapiro-Wilk test was utilized to assess the distribution of investigated features.

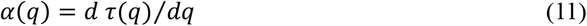

where, τ(*q*)=(*q*−1)*D*_*q*_ is known as the mass exponent

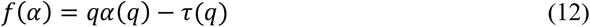

#### 2.3.2 Group Comparison

The Mann-Whitney U Test was applied to determine whether individual fractal parameters and the padding percentage exhibited significant differences between Normal and Malignant categories. Subsequently, Cliff’s delta test was conducted to assess the effect size of these differences.

Kruskal-Wallis Test was applied to evaluate the statistical differences across the four subtypes, and Dunn’s Post hoc Test with Bonferroni correction was performed to identify pairwise subgroup differences. The test was also performed for different percentile ranges (5^th^ and 95^th^, 1^st^ and 99^th^, 10^th^ and 90^th^, and 25^th^ and 75^th^), specifically for the succolarity reservoir parameter.

#### 2.3.3 Feature Interdependence and Principal Component Analysis (PCA)

Spearman’s Rank Correlation Coefficient was computed to analyze the relationships between features and between fractal features and subtypes. PCA was performed to investigate the capability of fractal features to capture distinct variance components in mammographic global textures and across subtypes.

#### 2.3.4 Classification Performance Evaluation

The Receiver Operating Characteristic (ROC) analysis was performed to assess the discriminative ability of individual fractal features.

The classification performance was also evaluated using supervised machine learning models, i.e., K-Nearest Neighbors (KNN) and Support Vector Machine (SVM), respectively. Stratified sampling was done to split the data into 80% training and 20% testing. A 5-fold stratified cross-validation was performed for optimizing the number of neighbors in the KNN model, while for SVM, it was done for optimization of the regularization parameter and kernel type (linear, Radial Basis Function (RBF), and polynomial). The selection of hyperparameters was done based on the Area Under the Curve (AUC) score, and the model was trained using the optimized parameters. The classification models were assessed based on the values of the Accuracy, AUC, F1-score, and Confusion Matrix metric on the test set.

An unsupervised machine learning approach, i.e., clustering analysis, was used to explore the subtypes-based natural groupings of mammogram images based on fractal features using the K-means clustering, Hierarchical clustering, and Density-based Spatial Clustering of Applications with Noise (DBSCAN) techniques. The quality of clusters was assessed by the Silhouette score metric.

### 3. Results

### 3.1 Differentiation between Normal and Malignant Categories

We evaluated the distributional properties of the computed fractal features across normal and malignant categories, and summary statistics (mean, standard deviation, minimum, and maximum) of each feature separately. Fractal dimension displayed a value of 2.634 ± 0.073 and 2.617 ± 0.089 for the malignant and normal cases, respectively, though a considerable overlap was observed in the distribution (Fig. 1(a)). Similarly, lacunarity showed significant overlapping between categories, with a value of 0.031 ± 0.018 and 0.033 ± 0.025 for the malignant and normal cases. Succolarity reservoir displayed a lower value for the malignant (0.008 ± 0.024) than the normal category (0.015 ± 0.041). It also displayed a higher maximum value (0.59) for the malignant cases as compared to the normal (0.51) as shown in Fig. 1(a). The Renyi dimensions (capacity, information, and correlation) displayed a similar trend between them in malignant and normal categories. The most pronounced difference between the malignant and normal categories was observed in multifractality strength (Fig. 1(a)) with corresponding values of 1.047 ± 0.215 and 0.973 ± 0.226, respectively. Fig. 1(b) presents the distribution of the investigated texture features for malignant and normal mammograms using box plots. The fractal dimension displayed a slightly higher median value, though, with a narrower interquartile range for the malignant cases compared to the normal. Lacunarity showed a similar distribution of median values with several outliers in both groups and substantial overlap between categories. A highly skewed distribution was observed for the succolarity reservoir parameter, with few extreme outliers and a slightly wider spread for the malignant cases compared to the normal ones. However, the median values were very low for both categories. The capacity dimension displayed little variation between categories, while the information and correlation dimensions remained relatively stable across samples with a small interquartile range. A comparable display of median values for the multifractality strength was observed for both categories, though the interquartile range shifted towards the higher values with a broader distribution for the malignant cases. Interestingly, all the features except lacunarity were found to display significant differences (*p* < 0.05) between normal and malignant mammograms from the Mann-Whitney U test. In addition, all the features were observed to follow a non-normal distribution from the Shapiro-Wilk test.

**Fig. 1.**
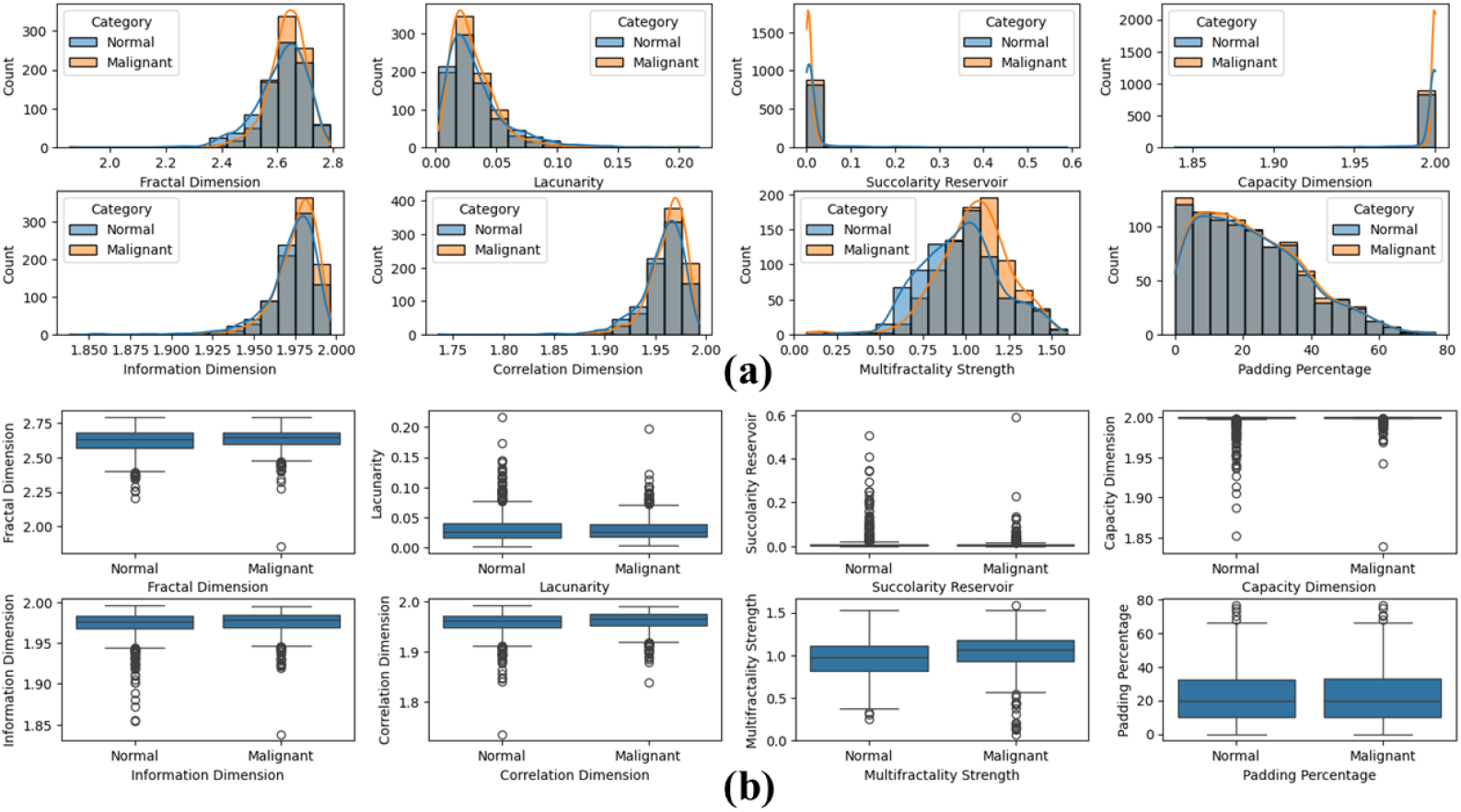
Graphical representation of **(a)** histograms and **(b)** box-plots for the analyzed fractal textural features corresponding to malignant and normal mammograms

We have also analyzed the padding percentage to avoid bias in interpreting results due to the concerned preprocessing technique. Nonetheless, the padding percentage remains almost identical across groups, as can be observed from both Fig. 1(a) and 1(b), individually. Also, the difference between categories was noted to be statistically non-significant from the Mann-Whitney U test. Subsequently, it was not considered as a covariate in further analyses.

The effect size of differences in the analyzed fractal features between normal and mammograms was quantified from the Cliff’s delta test. Subsequently, all the features except the multifractality strength showed a negligible effect size, while the latter displayed a small effect size. The computed values of Cliff’s delta are given in Table S1 (Supplementary Material).

Owing to the non-normal distribution of data, Spearman’s rank correlation test was performed to assess the presence of monotonic relationships and examine the interdependence between fractal-based texture features. The resulting correlation heatmap is shown in Fig. 2. The Fractal dimension demonstrated remarkable independence, exhibiting negligible correlations with all other parameters. In contrast, moderate and strong relationships were observed between several other parameters. The Renyi dimensions displayed exceptionally high intercorrelations, with information and correlation dimensions showing nearly perfect positive (= 0.99) and high-strength associations. Succolarity and capacity dimension exhibited a perfect negative correlation. Lacunarity showed a strong negative correlation with information and correlation dimensions while maintaining a moderate positive association with succolarity and multifractality strength. Multifractality strength demonstrated a moderate positive (negative) correlation with succolarity (capacity dimension).

**Fig. 2.**
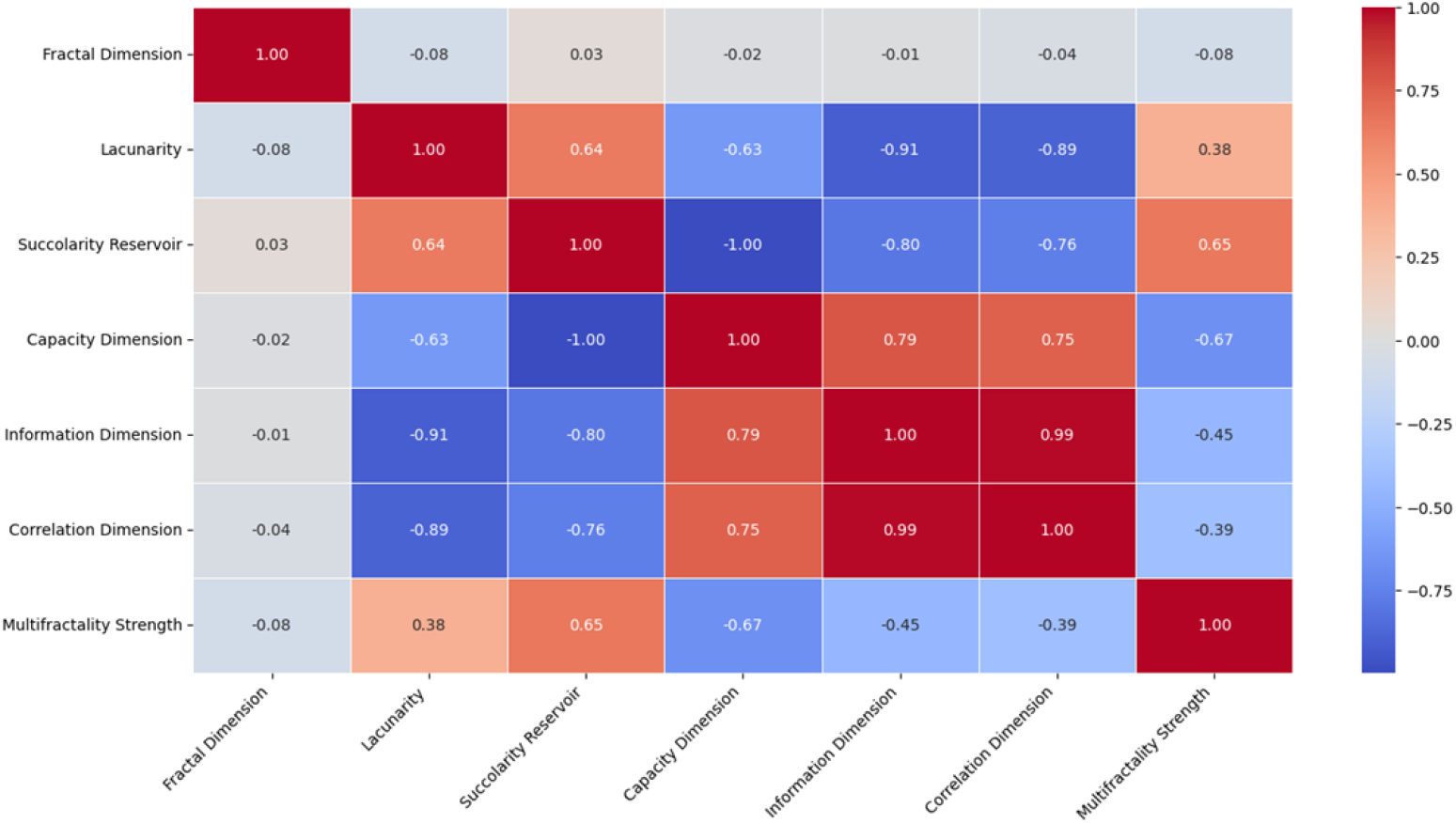
Heatmap representing the association between the investigated fractal features

The ROC analysis was performed to assess the classification efficacy of individual fractal-based texture features in distinguishing between malignant and normal mammograms. The AUC was computed for each feature to quantify its discriminative power. The results are displayed in Fig. 3. Among the features analyzed, the fractal dimension demonstrated the highest AUC, followed by multifractality strength and information dimension. The capacity dimension demonstrated moderate discriminative ability, while lacunarity and succolarity were less effective. The correlation dimension displayed the lowest AUC value. Also, the AUC value of 0.52 for the lacunarity feature indicates a performance close to random classification.

**Fig. 3.**
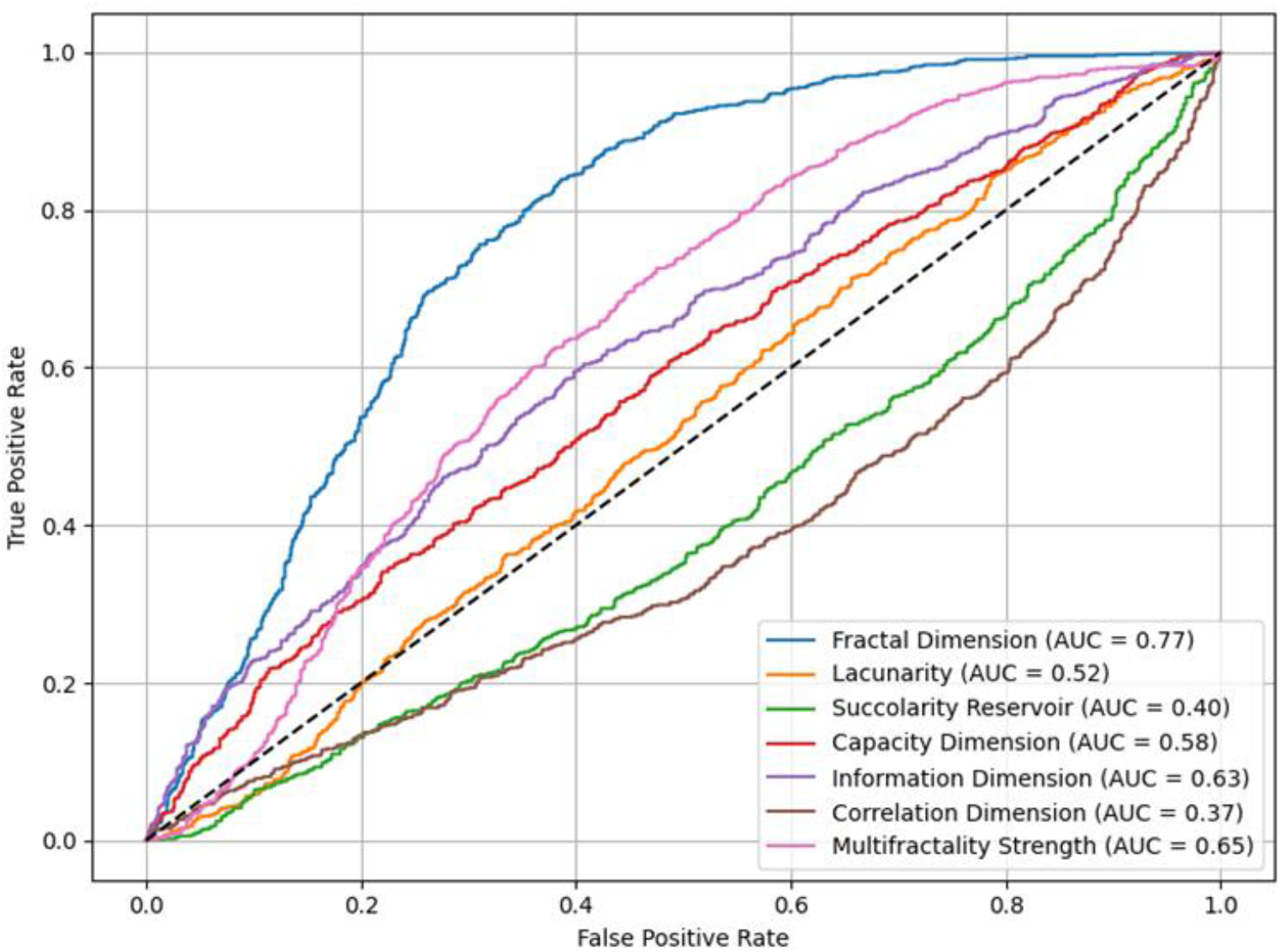
The Receiver Operating Characteristic curves corresponding to the investigated features

Principal Component Analysis was employed to evaluate the capability of fractal features in distinguishing between malignant and normal mammograms. The biplot (Fig. 4) illustrates the separation of categories based on the first two principal components. The first principal component (PC1) accounted for 58.14% of the total variance, while the second principal component (PC2) contributed 17.08%. In other words, the first two components capture 75.22% of the total variance in the data. Examination of component loadings revealed that PC1 was primarily characterized by large positive contributions from the Renyi dimensions with opposite negative contributions from lacunarity and succolarity. PC2 demonstrated strong positive loadings from multifractality strength, fractal dimension, and capacity dimension, with negative loading from the succolarity reservoir.

**Fig. 4.**
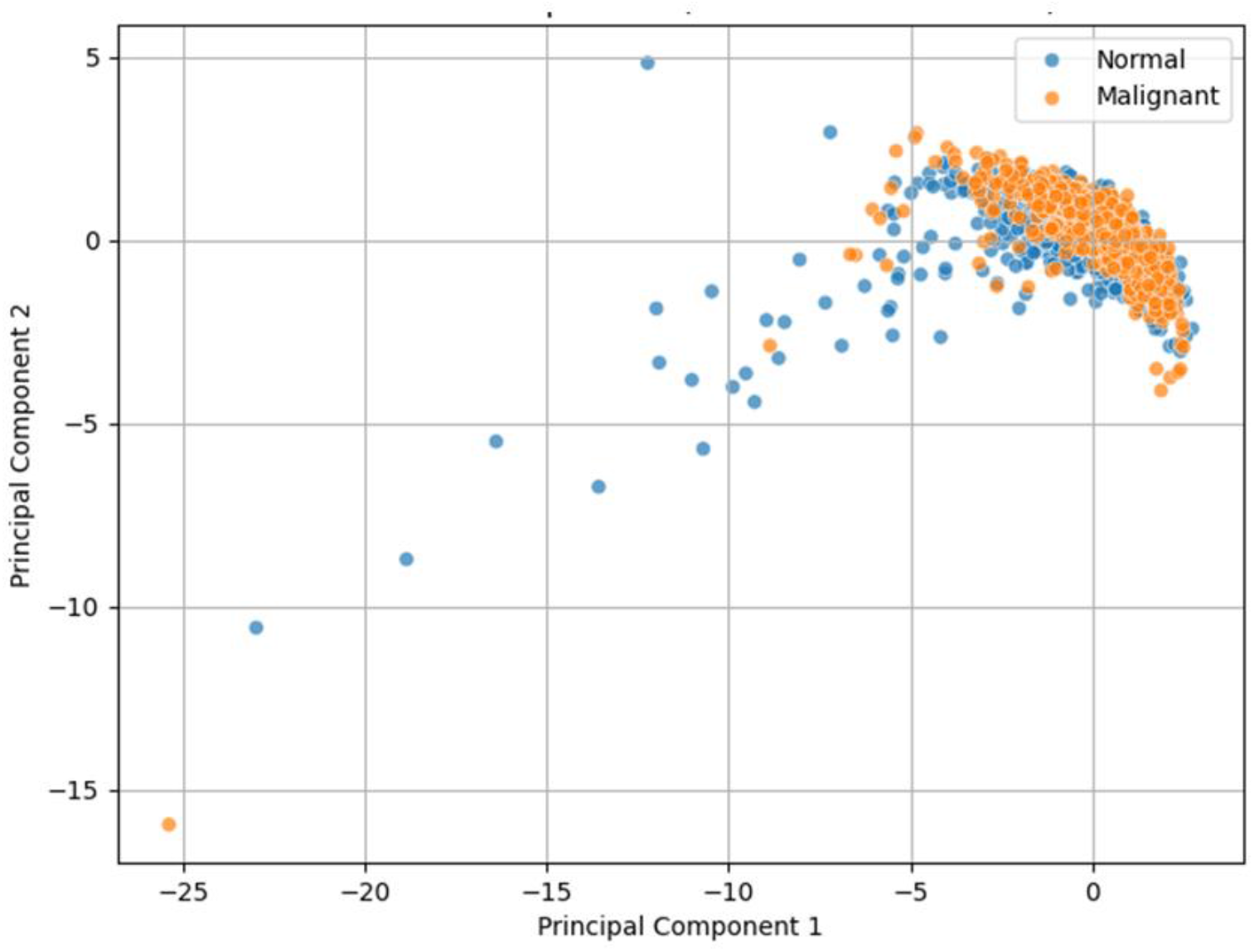
Biplot illustrating the separation of malignant and normal categories based on the first two principal components, taking into account 58.14% and 17.08% of variance in the dataset by Principal Component 1 and Principal Component 2, respectively

Nonetheless, a considerable overlap between malignant and normal categories was observed in Fig. 4, with malignant cases forming a more concentrated cluster primarily for the higher PC1 values, while normal cases showed greater dispersion with several extreme outliers at the lower negative end of PC1 and higher end of PC2.

Finally, to classify mammograms into malignant and normal categories, we employed a systematic machine learning approach using mono- and multi-fractal parameters as predictive features. We started with all the parameters; however, to improve model efficiency and interpretability, we implemented the Recursive Feature Elimination with a Random Forest Classifier as the base estimator. This ensemble-based feature selection method evaluates the importance of features by recursively eliminating the least significant ones while monitoring model performance. The analysis identified five optimal parameters, namely, fractal dimension, lacunarity, succolarity reservoir, capacity dimension, and multifractality strength. Using these selected features, we implemented and optimized the KNN and SVM classifiers. Subsequently, both optimized models achieved similar classification accuracy. The SVM model demonstrated superior discriminative capability with an AUC of 0.7253 compared to the KNN model’s AUC of 0.686. Confusion matrices revealed that both models showed a slight bias toward the malignant category, with high recall.

Nonetheless, to conceptually enhance the biophysical interpretability, we explored a dimensionally reduced model using only three parameters, i.e., fractal dimension, succolarity reservoir, and multifractality strength. The selection was based on their importance rankings from the previous statistical analyses. Here, the SVM model achieved notably strong performance with a higher accuracy than the KNN and the five-parameter SVM model. It also showed good discriminative ability with an AUC of 0.6526, an F1-score of 0.6917, high precision for normal cases, and strong recall for malignant cases. However, this model also showed a bias toward malignant classification as revealed by the confusion matrix. The category-specific performance metrics are shown in Table S2.

The resulting decision boundary of the SVM model is shown in Fig. 5. A complex, non-planar surface of the decision boundary underscores the non-linear relationship between fractal parameters and malignancy classification, with both malignant and normal cases, highlighted by red and blue, respectively, form partially overlapping clusters in the parameter space.

**Fig. 5.**
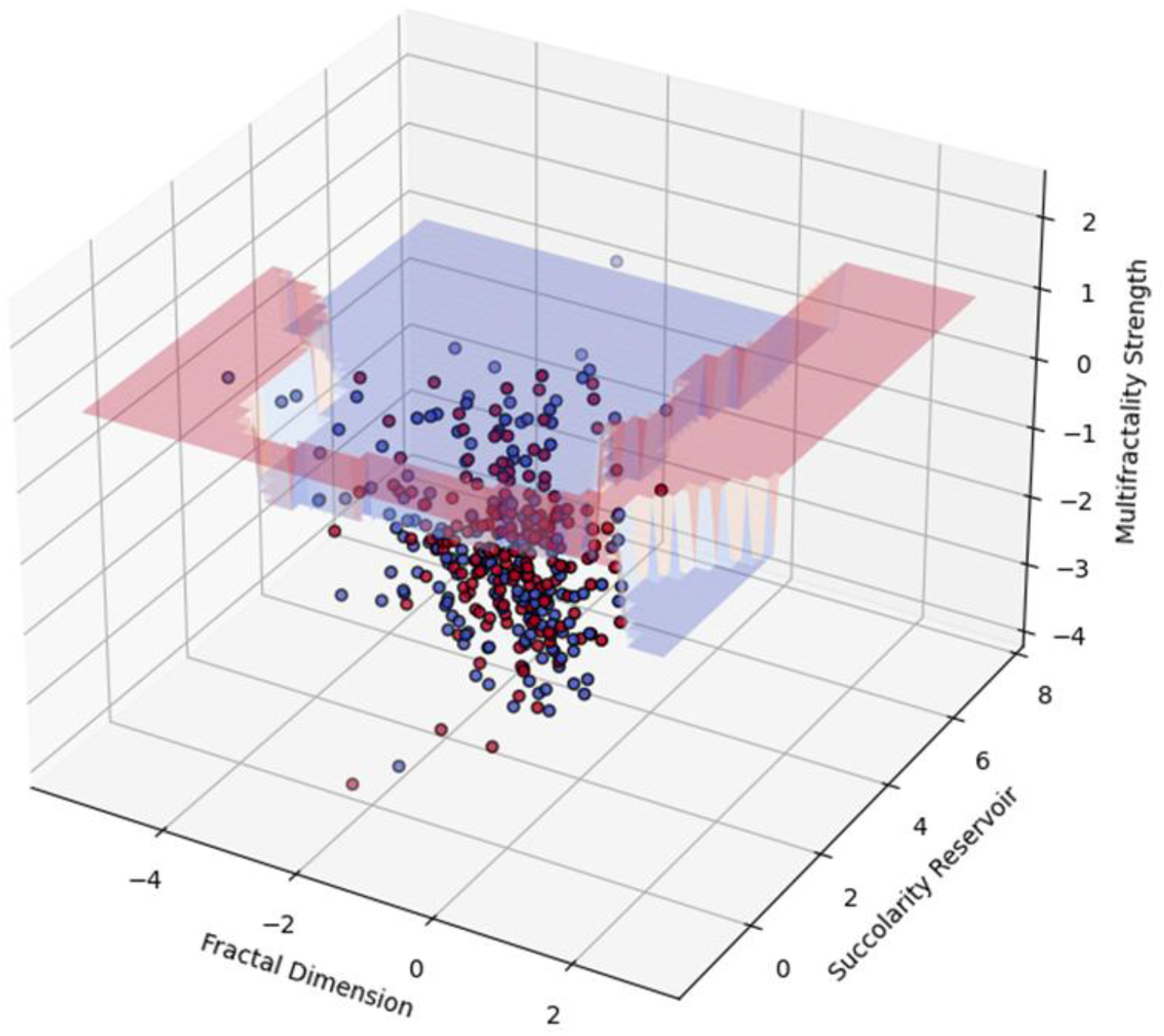
Pictorial representation of the three-dimensional decision boundary corresponding to the Support Vector Machine classification model. Here, the red and blue colored points represent the malignant and normal mammogram samples, and the boundaries define the regions where the classifier changes its prediction from malignant to normal or vice versa

### 3.2 Differentiation between Breast Carcinoma Subtypes

We studied the significance of the analyzed fractal textural features in molecular subtype differentiation. Fig. S1(a) displays the distribution characteristics of the features. The fractal dimension follows an approximately normal distribution with a slight negative skew, with a narrow peak indicating stable behavior across samples. In contrast, lacunarity exhibits a right-skewed distribution with a long tail, suggesting that while most samples have low values, a subset demonstrates significantly higher values. The succolarity reservoir, on the other hand, presents an extremely positive skew, with most values concentrated near zero. The Renyi dimensions display a similar distribution, characterized by high kurtosis and tight clustering 2.0. The multifractality strength follows an approximately normal distribution with moderate dispersion, with a small secondary peak at lower values, around 0.2-0.4. Violin plot (Fig. S1(b)) analysis further refined the understanding of these distributions across subtypes. The fractal dimension distributions are relatively symmetric across all subtypes, centered around 2.6-2.7, however, with greater variability in Luminal A and Luminal B. Lacunarity distribution confirms a positive skew across all subtypes, with the Luminal A cases showing higher density in the upper range. The succolarity reservoir distribution remains highly skewed, with Luminal B showing a slightly elevated body and a profuse upper region. Outliers in Luminal A and B subtypes, respectively, reach values up to 0.6, in contrast to lower maxima in HER2-enriched and triple-negative subtypes. The violin plots for the Renyi dimensions corroborated with the histogram results. Multifractality strength distributions remain approximately normal, with a more uniform spread of values for the Luminal B subtype.

The pairwise scatter plots in Fig. S2 provide additional insights into the interrelationship between fractal features. The diagonal plots confirm previously observed distributional differences across subtypes, particularly the broader distributions of fractal dimension and multifractality strength in Luminal B. Strong positive correlations are identified between information and correlation dimension, whereas a moderate negative correlation exists between lacunarity and fractal dimension. Additionally, multifractality strength and fractal dimension exhibit a moderate positive relationship. Outliers, particularly within Luminal B samples, appear consistently across multiple feature combinations, suggesting possible biological variations rather than measurement artifacts. Spearman correlation analysis reinforces these observations, demonstrating weak correlations between individual fractal features and subtypes, as shown in Fig. S3(a). Directional patterns indicate that the succolarity reservoir, information dimension, correlation dimension, and multifractality strength exhibit small positive correlations, while the fractal dimension, lacunarity, and capacity dimension show small negative correlations.

PCA analysis provides further evidence of the data structure and feature relationships. The clustering of points along PC1 suggests that this dimension captures the majority of the variance (54.69%) in the dataset, while PC2 accounts for 22.81% of the variance (Fig. S3(b)). However, despite dimensionality reduction, significant overlap between subtypes persists, confirming the absence of a monotonic relationship between fractal textural features and BC subtypes.

The above-mentioned highlights the non-discriminatory power of fractal features and was reinforced by the Kruskal-Wallis test, performed owing to the non-normal distribution of residuals, exhibiting non-significant differences for all the features across subtypes.

Nevertheless, based on the findings, which suggested the presence of outliers possibly influencing statistical comparisons, we conducted a more focused analysis by restricting the dataset to various percentile ranges and taking only the three significant parameters. The sample sizes varied from 159 (25-75 percentile) to 858 (1-99 percentile) cases. Kruskal-Wallis tests revealed statistically significant differences in succolarity reservoir across subtypes in three of the four percentile ranges-5^th^-95^th^ (*p* = 0.0393), 1^st^-99^th^ (*p* = 0.0181), and 10^th^-90^th^ (*p* = 0.0031). For the 25^th^-75^th^ percentile range, no significant difference was detected (*p* = 0.5896), possibly due to the substantially smaller sample size. Additionally, multifractality strength showed significant differences among subtypes in the 1^st^-99^th^ (*p* = 0.0147) and 10^th^-90^th^ (*p* = 0.0396) percentile ranges. The fractal dimension did not show any significant differences across subtypes in any of the percentile ranges. Post-hoc analysis using Dunn’s test for succolarity reservoir revealed that the most consistent pairwise differences were observed between Luminal A and Luminal B with (*p* = 0.099, 0.068, 0.023) in the 5^th^-95^th^, 1^st^-99^th^, and 10^th^-90^th^ range, respectively, as well as between Luminal B and triple-negative subtypes (*p* = 0.023) in the 10^th^-90^th^ range.

Spearman correlation analyses between fractal textural features and BC subtypes did not yield statistically significant associations across any of the percentile ranges analyzed. PCA revealed consistent patterns across 5^th^-95^th^, 1^st^-99^th^, and 10^th^-90^th^ range, with the succolarity reservoir having the highest contribution to PC1. For the 25^th^-75^th^ range, multifractality strength showed the highest loadings to PC1. Notably, fractal dimension consistently dominated PC2 across all percentile ranges.

Finally, we employed multiple unsupervised clustering algorithms across the selected percentile ranges to identify natural groupings in the data based on fractal parameters. K-Means clustering yielded the highest silhouette scores across all percentile ranges (0.271-0.305), indicating better-defined clusters compared to hierarchical (0.234-0.245) and DBSCAN (-0.052-0.077) clustering. DBSCAN identified varying numbers of clusters (noise points), i.e., 4 (155), 3 (193), 9 (117), and 2 (27) in the 5^th^-95^th^, 1^st^-99^th^, 10^th^-90^th^, and 25^th^-75^th^ range, respectively. Overall, the analysis demonstrated that while there are discernible patterns in the analyzed features across subtypes, the clustering structures are not strongly defined, as evidenced by the moderate silhouette scores across all algorithms and percentile ranges. The associated pictorial representations of K-Means, Hierarchical, and DBSCAN clustering analyses results are shown in S4(a), S4(b), and S4(c), individually.

## 4. Discussion

This study aimed to investigate the efficacy of fractal-based global parenchymal texture analysis of ROIs in differentiating between normal and malignant breast tissues in mammograms, with implications regarding insights for BC research and non-invasive diagnosis/detection. Daye *et. al*. (47) speculated that parenchymal textural patterns provide complementary information, along with mammographic density, symptomatic of malignancy. Our findings highlight the potential of model-based approaches, particularly mono- and multi-fractal analysis, in capturing the spatially complex structural and connectivity patterns in breast tissues that may be indicative of malignancy. Among the investigated textural features, fractal dimension, multifractality strength, and succolarity reservoir emerged as the most significant features for distinguishing normal from malignant tissues.

### 4.1 Implications for BC Detection or Diagnosis

The supervised machine learning approaches demonstrate that fractal-based texture features can moderately differentiate between malignant and normal mammograms. However, the reduced SVM model using three features (fractal dimension, multifractality strength, and succolarity reservoir) achieved comparable or better performance than models using the complete feature set, highlighting the critical role of these fractal measures in mammographic texture characterization. Nonetheless, the non-linear decision boundary observed in the SVM model (Fig. 5) suggests that the relationship between fractal parameters and malignancy is complex and multi-faceted, supporting the notion that cancer progression involves multiple, interdependent structural alterations that cannot be captured by a single measure.

The findings also indicate that the succolarity reservoir provides complementary detection or diagnostic information, despite its lower standalone discriminative power.

#### 4.1.1 Fractal Dimension

The fractal dimension demonstrated the highest discriminative power, as determined by ROC analysis, among all the investigated features. This metric quantifies the complexity, intricacy, irregularity, and space-filling capacity of breast textural patterns in mammographic images. However, these attributes have distinct interpretations-complexity implies systems or patterns composed of many interconnected elements, intricacy implies elaborate detail, fine structure, and a high level of internal differentiation, irregularity implies deviation from symmetrical or predictable patterns, and space-filling capacity implies the ability of structures or patterns to effectively occupies the available space within its domain. From a biophysical perspective, treating fractal dimension as a measure of complexity and space-filling capacity is more appropriate from a biological viewpoint since it can provide insights into tumor architecture and tissue organization. In addition, it could serve as a geometrical analogue to the variation or disruption in the tissue density, structural distribution, and architectural patterns during oncogenesis. An increased fractal dimension in malignant tissues reflects irregular space-filling growth patterns, subsequently, a more complex architectural organization of tissues. The phenomenon reflects the underlying cellular and tissue heterogeneity and altered stromal organization, which are hallmarks of tumor microenvironments. Tumor progression is driven by processes like unregulated cell proliferation and extracellular matrix remodeling, all of which contribute to more complex tissue structures (48). Nonetheless, in the context of parenchymal texture, the fractal dimension serves as a quantitative measure of tissue heterogeneity. Our findings align with existing literature demonstrating a positive correlation between high parenchymal complexity and increased malignancy risk (49). Parenchymal complexity highlights the textural variation in breast tissue patterns, which in turn is influenced by the composition, spatial organization, and structural distribution of cellular and extracellular matrix within tissues. In this regard, the higher fractal dimension in malignant cases can correspond to disruption of tissue homeostasis with high variability in epithelial and stromal structures. Also, since dense breasts are primarily composed of fibroglandular tissues (49), a higher fractal dimension corresponds to an augmentation in texture complexity.

The relative independence of this feature from the others, as indicated by the Spearman correlation analysis, also supports its unique role as a descriptor of tissue global complexity.

Finally, the fractal dimension has been reported to be a measure of self-similarity in textural patterns (49); however, parenchymal patterns can manifest as either directional-independent or dependent variations in texture. In this regard, the self-similarity should be clearly described or interpreted as either statistical or self-affine. In our study, the fractal dimension would signify self-affinity since it is interpreted in terms of grey-scale or *z* − *axis* variations. Nonetheless, the notion of self-similarity is not very visually obvious, especially in the case of the breast.

#### 4.1.2 Multifractality Strength

Multifractality strength exhibited the 2^nd^-highest discriminative ability from ROC analysis but also showed a small but notable effect size (Cliff’s delta test) in differentiating between normal and malignant breast tissues. Its higher value for malignant mammograms compared to normal cases suggests that tumor textures exhibit greater variability in scaling properties across multiple spatial scales. The results corroborate with findings from Joseph and Pournami (28), who demonstrated that multifractal analysis effectively captures the heterogeneous nature of malignant tissues, distinguishing them from normal ones. The observation was also reinforced by the negative value of this measure in Cliff’s delta test, suggesting its augmented values in malignant tumors. Unlike the fractal dimension metric, which provides a single measure of scaling, multifractality strength accounts for variations in scaling behavior, consequently, textural complexity at different resolutions.

In the context of grey-scale mammogram images, malignant transformation involves diverse cellular populations with varying proliferative and invasive capacities, creating heterogeneous tissue densities that appear as regions of varying pixel intensities, manifesting as increased multifractality strength in the distribution of grey-scale values across the analyzed images. Notably, despite the limitation of mammography, where textural features reflect an admixture of overlapping fatty, fibroglandular, and superficial skin tissues (49), limiting the observation of parenchymal textural patterns, the observed increase in multifractality strength for the malignant category suggests that tumor-associated heterogeneity remains detectable. Moreover, as BC is predominantly associated with fibroglandular tissue characteristics (49), the ability of this measure to differentiate malignant from normal mammograms highlights its significance in quantifying malignancy-driven alterations in tissue complexity, even in the presence of overlapping structures.

#### 4.1.3 Succolarity Reservoir

The succolarity reservoir parameter, which quantifies the global percolation capacity or latent connectivity of breast tissues, was found to be significantly lower in the malignant category compared to the normal category. This behavior was reinforced by the delta test, which yielded a positive value for this metric.

From a biophysical standpoint, the succolarity reservoir can be defined as the fraction of percolative pathways (represented by black pixels) relative to the total tissue area, with white pixels representing structural barriers and grey pixels indicating latent connectivity. This metric provides a global, direction-independent measure of textural percolation in breast tissues.

In normal breast parenchyma, particularly in fibroglandular tissues, this measure is expected to be higher owing to the presence of well-connected ductal and stromal structures that facilitate percolation-like continuity. This conjecture is supported by the differential behavior of the succolarity parameter between normal, benign, and malignant mammograms (50). It can be hypothesized that the organized tissue architectural patterns with hierarchical branching and ductal networks provide a regular spatial relationship between epithelial and stromal components, contributing to greater connectivity pathways. Patel *et al*. (51) studied the global breast tissue stiffness using magnetic resonance elastography and reported that normal tissues maintain an organized stromal framework with compliant mechanical properties. The observation is in agreement with our observation as it implies higher tissue connectivity. In contrast, malignant transformation disrupts connectivity through processes like augmented fibrosis and disordered cellular arrangements facilitated by tumor expansion. It has been testified that mammographically dense tissues exhibit varying levels of collagen density, stroma stiffness, and epithelial cell density, and are found to be related to the progression of BC (51–53). Thus, it can be speculated that altered collagen deposition and alignment, and disruption of the ductal-stromal spatial relationship result in reduced connectivity patterns. These changes could manifest themselves as more whitish or fewer greyish pixels in mammogram images, thereby lowering the value of this feature in the malignant category.

In addition, the grey pixels represent areas that could become percolated if the structural barriers were breached. In normal mammograms, grey regions may indicate latent connectivity through ductal and stromal bridges; however, in malignancy, these potential connections are lost due to the remodeling of the extracellular matrix and tumor-induced architectural disruption, including disruption of basement membrane continuity and stromal reorganization. The perfect negative correlation between the succolarity reservoir and capacity dimension (Fig. 2) suggests that these features capture opposite aspects of tissue architecture. In other words, as tissue structural complexity increases (higher capacity dimension), the connectivity decreases. This is consistent with the notion that malignant transformation introduces fine-scale complexity but disrupts large-scale organization.

Given its ability to reflect global connectivity patterns, the succolarity reservoir may serve as a descriptor for the parenchymal organization in breast tissue analysis. Its potential role in breast density classification, malignancy risk assessment, percolation-based tumor microenvironment, and tissue mechanical properties characterization warrants further exploration.

### 4.2 Molecular Subtype Differentiation

The analysis of fractal textural features across BC molecular subtypes revealed limited discriminative power when considering the entire dataset. However, the focused analysis with different percentile ranges revealed that the succolarity reservoir was the most discriminative factor across the subtypes, showing statistical significance in three of the four examined percentile ranges.

The differentiation between Luminal A and Luminal B subtypes (*p* = 0.023) in the 10^th^-90^th^ range, suggesting that these molecularly distinct entities also exhibit differences in the spatial organization and connectivity patterns of breast tissues. Luminal B tumors, characterized by high proliferation rates and more aggressive behavior compared to Luminal A (35,54), exhibited enhanced latent-connectivity as evidenced by their higher median succolarity reservoir value. The results qualitatively corroborate the observed correlation between radiomics signatures and BC molecular subtypes, as reported by Leithner *et al*. (55). This finding suggests that, despite their more aggressive nature, Luminal B tumors may maintain or increase certain aspects of tissue connectivity. The higher values of this feature in the Luminal B subtype might also relate to differences in angiogenesis patterns, stromal organization, or cellular arrangement that can facilitate increased percolation capacity despite the more aggressive growth behavior of the tissues. The observation highlights the complex relationship between molecular subtypes and their physical manifestation in tissue architecture.

Similarly, the significant difference between Luminal B and triple-negative subtypes is visible in the distribution patterns, with Luminal B consistently showing higher values for the succolarity reservoir parameter. This finding suggests that triple-negative BC, despite its known aggressive growth patterns (54), may exhibit reduced tissue connectivity compared to Luminal B. The triple-negative subtype is known to display a circumscribed, round shape rather than a spiculated appearance (56,57). In addition, they have been reported to possess the highest tumor roundness score among subtypes with rim enhancement (58–60). Finally, triple-negative cases were reported to exhibit larger masses and more heterogeneous texture from radiomics analysis (61,62). These architectural distinctions can be correlated to a more compact yet aggressive and heterogeneous tumor growth, with distinct boundaries, contributing to a decrease in tissue latent-connectivity.

While the fractal dimension parameter showed no significant differences across subtypes, its relatively stable contribution to PC2 across all percentile ranges suggests that global structural complexity is an orthogonal aspect of breast tissue architecture that is less subtype-specific. The emergence of multifractality strength as the most influential feature in the 25^th^-75^th^ percentile range is intriguing however, no statistically significant difference was found between subtypes. This observation implies that, similar to global structural complexity, variability in scaling properties across multiple intratumoral spatial scales is less subtype-specific.

The modest silhouette scores across clustering algorithms indicate that fractal-based features provide meaningful discrimination, but they do not completely recapitulate established molecular classifications. These scores indicate that textural heterogeneity represents a distinct dimension of tumor biology that complements genomics and proteomics signatures.

### 4.3 Methodological Considerations and Limitations

Several methodological considerations are critical for interpreting our findings. First, to ensure robust comparisons, we standardized the ROI dimensions through aspect ratio-preserving resizing with neutral grey padding. Importantly, the statistical non-significance of padding differences across categories confirms that our preprocessing approach did not introduce systematic bias. This ensures that the observed variations in fractal features reflect genuine differences in tissue texture, subsequently, architecture rather than artifacts of image processing. Additionally, by analyzing the entire ROIs rather than specific lesions using raster boxes, our approach captures global textural differences between normal and malignant tissues. This strategy is particularly valuable in cases where lesions lack well-defined boundaries, a common challenge in mammography interpretation; however, significantly important for early detection/diagnosis. However, it also implies that small, localized malignant regions may be underrepresented in the analysis. Further studies may benefit from hybrid approaches, like combining global textural features with localized ones using raster and sliding box methods.

One notable limitation is that the succolarity reservoir is highly sensitive to spatial connectivity patterns and pixel variations, which may vary with tumor size, breast density, and imaging resolution. While our findings suggest that this feature is reduced in malignant tissues, future work should investigate its robustness across different imaging modalities, e.g., computed tomography and magnetic resonance imaging. Additionally, the observed perfect negative correlation between this feature and capacity dimension suggests a trade-off between tissue textural complexity and global connectivity patterns. This observation poses an important question i.e., to what extent does increasing structural complexity inherently reduce connectivity properties? The relationship warrants further exploration using computational modeling and *in vitro* tissue simulation.

Finally, while global texture analysis using fractal parameters demonstrated promising detection/diagnostic performance, the overlap between malignant and normal categories in PCA indicates that fractal features do not capture all relevant differences. This indicates that integrating other imaging processing and analysis techniques, such as contrast enhancement patterns, sliding-box algorithm, and transform-based methods, could improve detection or diagnostic precision. Another key observation is that our KNN and SVM models exhibited a slight bias toward malignant classification. While this may increase false-positive rates, it aligns with clinical priorities, i.e., minimizing false negatives is more critical than avoiding false positives. Nonetheless, future models should explore calibrating decision thresholds to optimize specificity while maintaining high sensitivity.

### 4.4 Comparison with Previous Studies

Our findings align with previous reports about higher fractal dimensions in malignant lesions (5,31–34). However, most prior studies have focused on tumor boundary fractal analysis, whereas our results emphasize the significance of global textural complexity in capturing malignancy-related architectural alterations. Our conceptual approach holds significance for early detection/diagnosis, as well-defined boundaries are a signature of high-grade tumors (57). Also, the significance of multifractality strength in differentiating breast tissue types builds upon previous research suggesting that multifractal analysis enhances the characterization of tumor heterogeneity in mammograms (28). This study provides further evidence that multi-fractality strength captures non-trivial, multi-scale variations associated with tumor microstructural, subsequently, tissue’s architectural complexity.

While previous studies have not extensively explored percolation-based measures in breast imaging, our results suggest that the succolarity reservoir provides valuable insights into tissue connectivity and architectural and organizational degradation in cancerous transformations. Further studies could further explore this parameter’s role in predicting tissue’s mechanical properties and tumor progression, and treatment response.

### 4.5 Future Directions

While this study provides a proof-of-concept, an integration of fractal-based texture analysis with other imaging features, such as morphological characteristics or with statistics-based/ transform-based methods, could improve diagnostic accuracy. The application of deep-learning approaches to extract and interpret fractal features might uncover more subtle patterns that differentiate malignant from normal tissues or distinguish between subtypes.

Significant differences in succolarity reservoir features across subtypes in specific percentile ranges suggest the potential for developing more targeted texture analysis approaches. Further studies could focus on identifying optimal ranges or regions for texture analysis that maximize subtype differentiation. Additionally, longitudinal studies examining changes in fractal features during cancer progression or treatment response could provide valuable insights into the dynamics of tissue architecture in BC. Such studies might elucidate how radiographic texture variations impinge on functionally different significant processes in breast carcinoma.

## 5. Conclusion

The performed study demonstrates the significance of fractal-based global parenchymal texture analysis in the differential diagnosis between malignant and normal tissues in mammograms. Fractal dimension, multifractality strength, and succolarity reservoir emerged as the most significant features, each capturing distinct aspects of tissue architecture that reflect underlying differences between malignant and normal tissues. While these parameters exhibited a limited ability to differentiate between molecular subtypes across the entire dataset, focused analyses revealed potential for subtype differentiation with specific data ranges. The findings contribute to the growing body of evidence supporting the use of texture analysis in mammographic interpretation and highlight the potential of model-based approaches in capturing complex spatial patterns in breast tissues. Further integration of these approaches with other imaging features and molecular-scale data could enhance our understanding of the radiographic-pathological correlation in BC and improve non-invasive diagnostic accuracy.

## Acknowledgment

**A.D**. acknowledges the Department of Biotechnology (DBT)-India for the Research Associate fellowship vide Award Letter No. DBT-RA/2023/January/NE/3594. **A.D**. also acknowledges Prof. Helmut Ahammer, Division of Medical Physics and Biophysics, Medical University of Graz, Graz, Austria, for the guidance regarding batch computation of fractal parameters.

**R. B**. acknowledges support from the Indo-French Centre for the Promotion of Advanced Research (69T08-2).

**M. K. J**. acknowledges support from the Param Hansa Philanthropies.

