## Supplementary Material for "Fractal Measures as Predictors of Histopathological Complexity in Breast Carcinoma Mammograms"

**Table S1** Tabulated result of the Cliff's delta test

| Parameter | Delta Value |
|---|---|
| Fractal Dimension | -0.101 |
| Lacunarity | 0.005 |
| Succolarity Reservoir | 0.128 |
| Capacity Dimension | -0.127 |
| Information Dimension | -0.112 |
| Correlation Dimension | -0.143 |
| Multifractality Strength | -0.22 |

**Table S2** Listed value of metrics highlighting the classification performance of the supervised machine learning models

| Model | Accuracy | AUC | F1-Score |
|---|---|---|---|
| KNN | 0.6264 | 0.6262 | 0.6356 |
| SVM | 0.6545 | 0.6526 | 0.6917 |

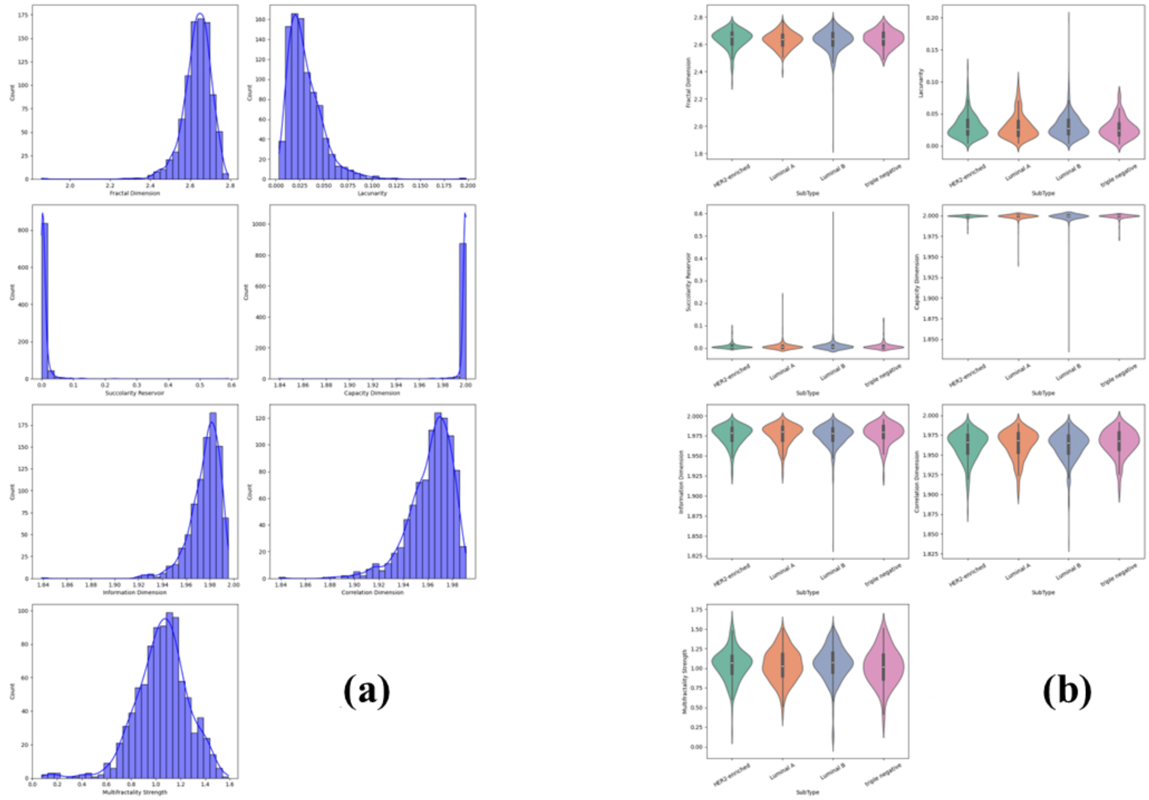


**Fig. S1** Graphical representation of **(a)** histograms and **(b)** violin plots for the analyzed fractal textural features corresponding to breast carcinoma molecular subtypes

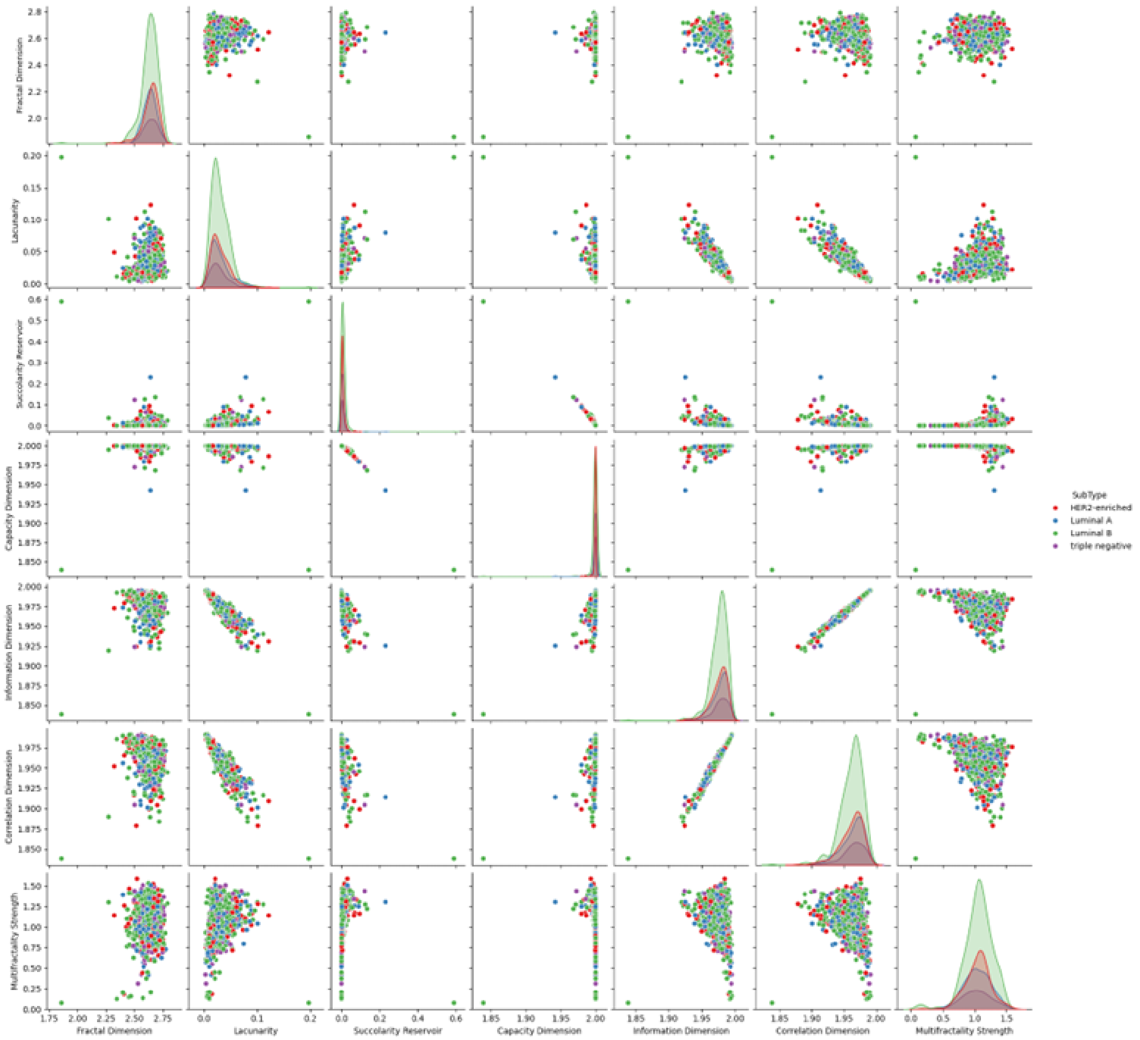


**Fig. S2** Pictorial representation of the interrelationship between the fractal features across subtypes

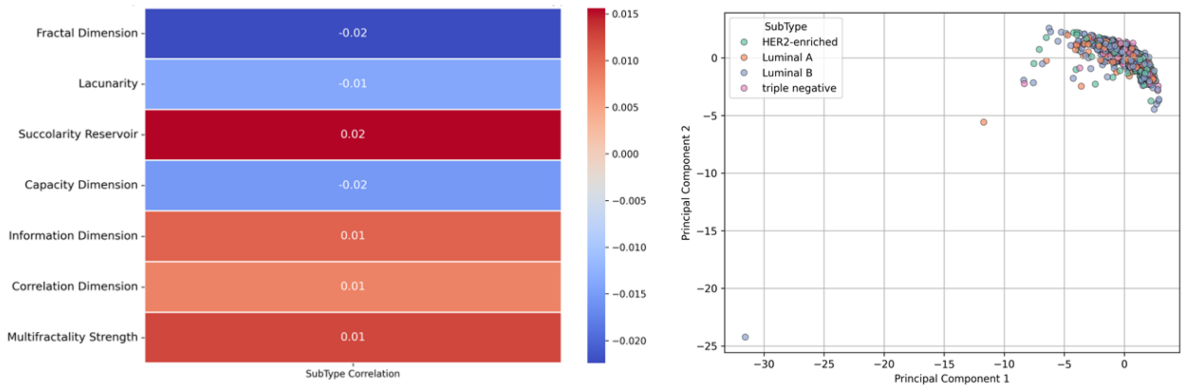


**(a)** **(b)**

**Fig. S3 (a)** Heatmap representing the association between features across subtypes and **(b)** Biplot illustrating the separation of molecular subtypes based on the first two principal components

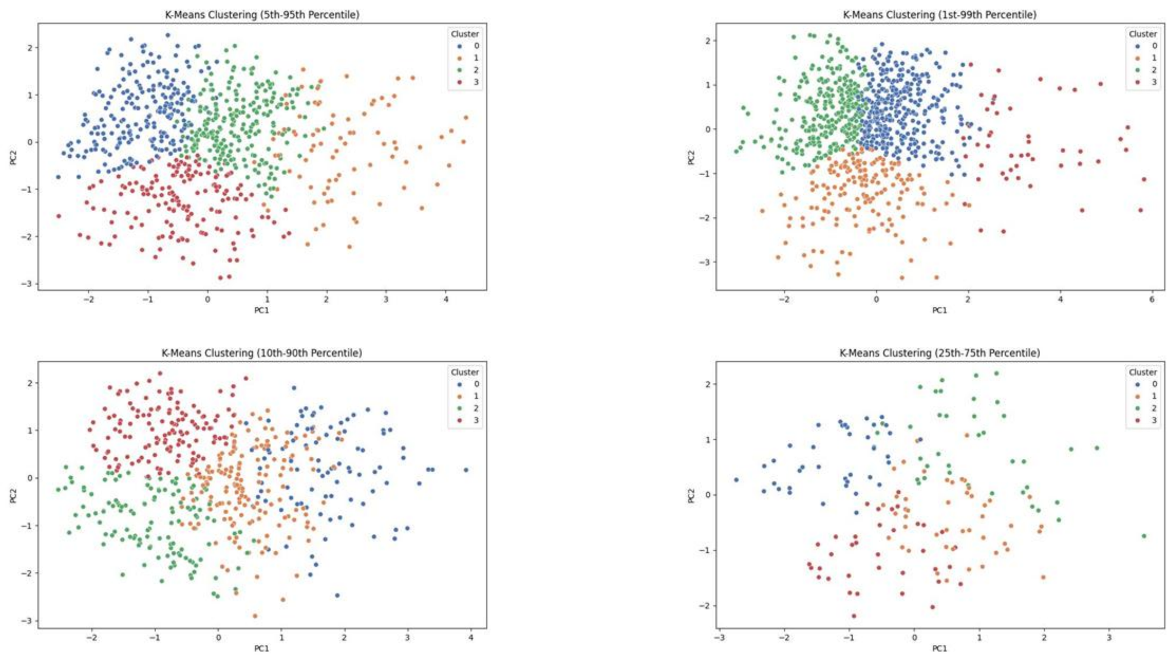


**Fig. S4(a)** Graphic illustration of the natural groupings in the subtype data based on fractal parameters using K-Means clustering

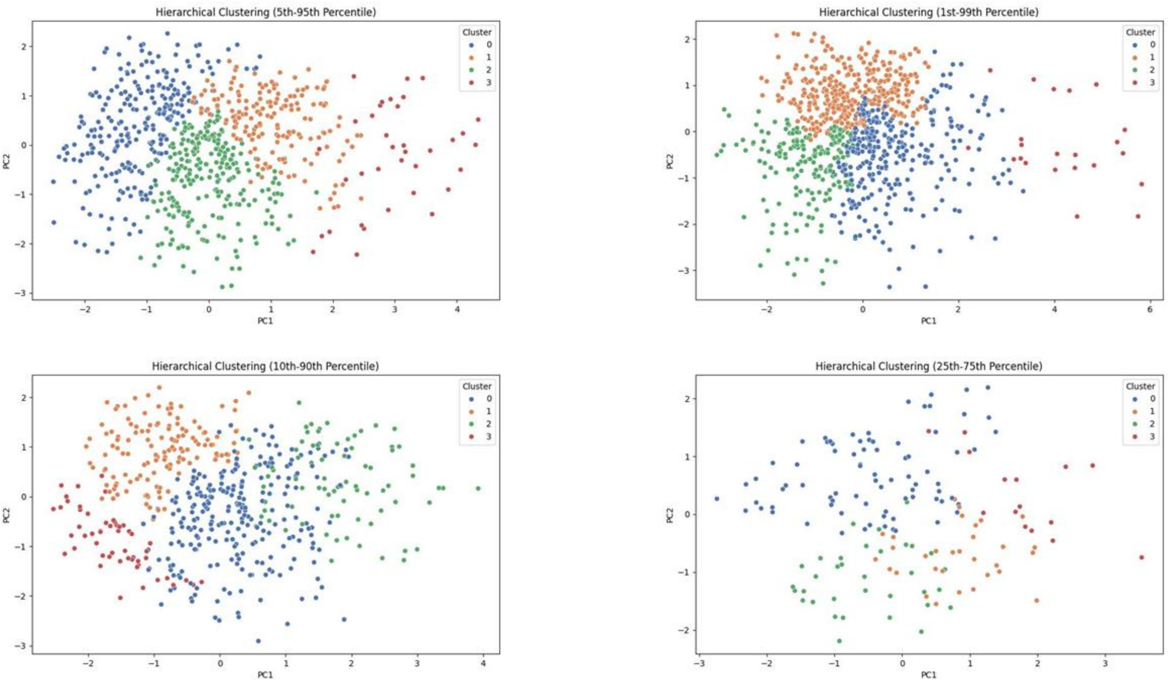


**Fig. S4(b)** Graphical representation of the natural groupings in the subtype data based on fractal parameters using Hierarchical clustering

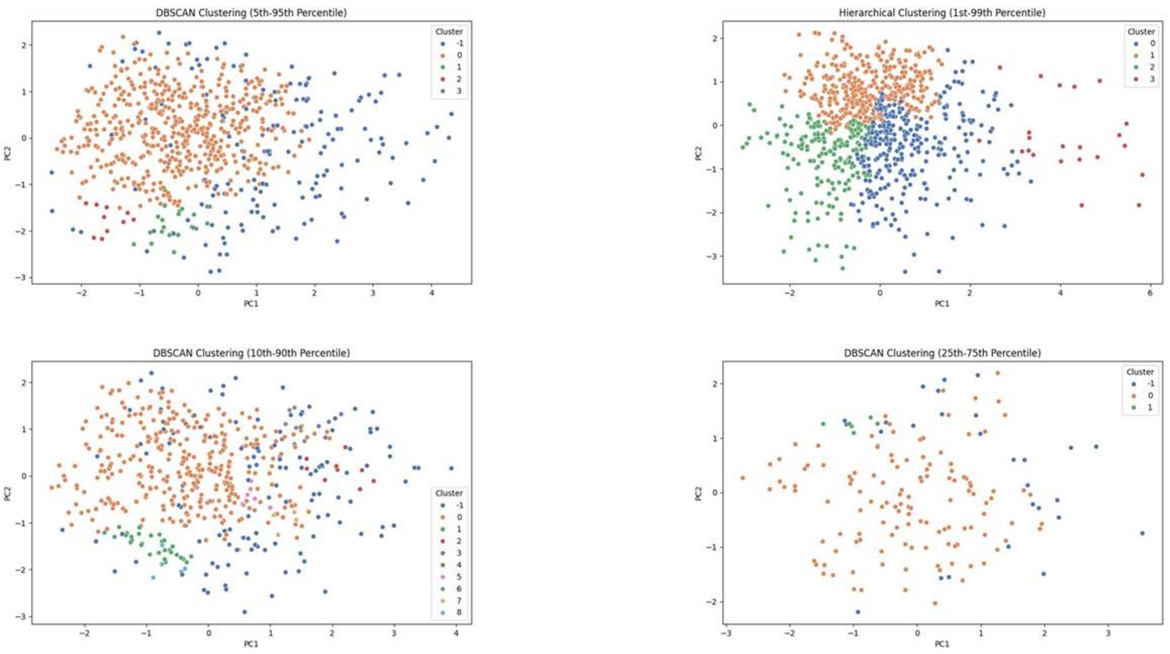


**Fig. S4(c)** Graphical representation of the natural groupings in the subtype data based on fractal parameters using Density-based Spatial Clustering of Applications with Noise (DBSCAN) clustering
